# Anti-frameshifting ligand active against SARS coronavirus-2 is resistant to natural mutations of the frameshift-stimulatory pseudoknot

**DOI:** 10.1101/2020.06.29.178707

**Authors:** Krishna Neupane, Sneha Munshi, Meng Zhao, Dustin B. Ritchie, Sandaru M. Ileperuma, Michael T. Woodside

**Affiliations:** Department of Physics, University of Alberta, Edmonton AB T6G 2E1, Canada

## Abstract

The coronavirus SARS-CoV-2 causing the COVID-19 pandemic uses −1 programmed ribosomal frameshifting (−1 PRF) to control the expression levels of key viral proteins. Because modulating −1 PRF can attenuate viral propagation, ligands binding to the viral RNA pseudoknot that stimulates −1 PRF may prove useful as therapeutics. Mutations in the pseudoknot have been observed over the course of the pandemic, but how they affect −1 PRF and the activity of inhibitors is unknown. Cataloguing natural mutations in all parts of the SARS-CoV-2 pseudoknot, we studied a panel of 6 mutations in key structural regions. Most mutations left the −1 PRF efficiency unchanged, even when base-pairing was disrupted, but one led to a remarkable three-fold decrease, suggesting that SARS-CoV-2 propagation may be less sensitive to modulation of −1 PRF efficiency than some other viruses. Examining the effects of one of the few small-molecule ligands known to suppress −1 PRF significantly in SARS-CoV, we found that it did so by similar amounts in all SARS-CoV-2 mutants tested, regardless of the basal −1 PRF efficiency, indicating that the activity of anti-frameshifting ligands can be resistant to natural pseudoknot mutations. These results have important implications for therapeutic strategies targeting SARS-CoV-2 through modulation of −1 PRF.

---

The severe acute respiratory syndrome coronavirus-2 (SARS-CoV-2) causing the COVID-19 pandemic that is currently sweeping the globe features a −1 programmed ribosomal frameshifting (−1 PRF) site.^1^ −1 PRF, which involves a shift in the reading frame of the ribosome at a specific location in the RNA message to generate an alternate gene product, is stimulated by a structure in the mRNA—typically a pseudoknot—that is located 5–7 nt downstream of the ‘slippery’ sequence where the reading-frame shift occurs.^2,3^ The −1 PRF signal is essential to SARS-CoV-2: the frameshift gene products include the RNA-dependent RNA polymerase that is required for viral replication. Previous work on SARS-CoV showed that mutations suppressing −1 PRF significantly attenuated viral propagation in cell culture, by up to several orders of magnitude.^4–6^ Indeed, work on other viruses has found more generally that −1 PRF levels must typically be held within a relatively narrow range to avoid attenuation.^7,8^ As a result, frameshift-stimulatory structures are potential targets for anti-viral drugs.^9–18^

The pseudoknot stimulating −1 PRF in SARS-CoV-2 has a 3-stem architecture that is characteristic of coronaviruses, in contrast to the more common 2-stem architecture of most frameshift-stimulatory pseudoknots in viruses.^19^ Although the pseudoknot sequence is highly conserved among coronaviruses,^20^ several mutations have been identified in the pseudoknot from viral samples isolated from COVID-19 patients during the pandemic, tracked in the GISAID database.^21^ Some of these mutations have been seen in only single patients; others have been found in patients from many different regions of the world (Fig. 1, right). Understanding how these mutations affect −1 PRF in SARS-CoV-2 can provide insight into issues of central relevance to developing therapeutic −1 PRF modulators. Such issues include the range of −1 PRF levels over which the virus can survive and propagate (defining the degree of modulation needed to achieve a therapeutic effect), the regions of the pseudoknot that are most sensitive to disruption (and hence most profitable to target therapeutically), and the extent to which mutations in the pseudoknot may be able to induce drug resistance. However, the effects of naturally occurring mutations in the SARS-CoV-2 pseudoknot on the levels of −1 PRF in SARS-CoV-2 and any changes they may induce in the activity of anti-frameshifting ligands have not yet been investigated. Here we do so, focusing on a ligand previously shown to inhibit −1 PRF in SARS-CoV.

**Fig. 1:**
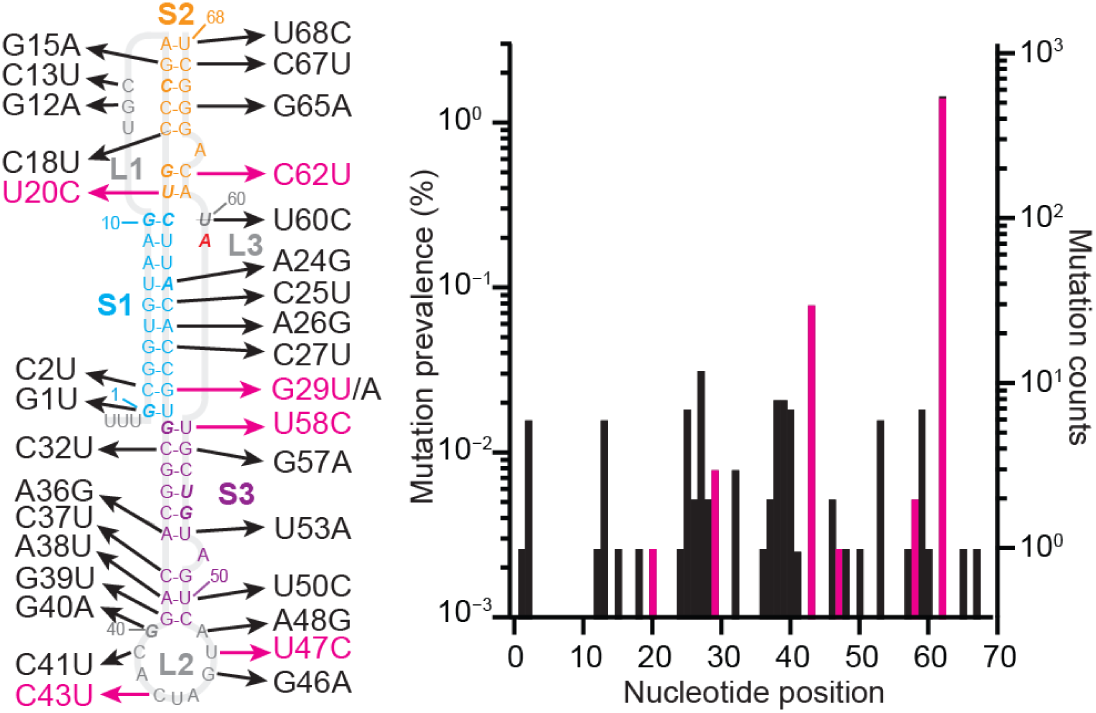
Natural mutations in SARS-CoV-2 pseudoknot. Left: Mutations identified from COVID-19 patients occur in all regions of the pseudoknot structure. Bases shown in italic are protected against nuclease digestion.^6^ Right: Most mutations have low occurrence. Mutations studied herein shown in magenta.

Surveying over 40,000 patient-derived sequences deposited in the GISAID database as of June 9, 2020 to identify mutations in the frameshift-stimulatory pseudoknot, we found that ∼1.8% of all sequences contained mutations, affecting 33 of the 68 nucleotides in the pseudoknot. In all cases but one, mutations involved single-nucleotide polymorphisms, without insertions or deletions; the exception involved the mutation of 3 adjacent nucleotides (A38–G40). These mutations occurred in all regions of the secondary structure (Fig. 1), distributed relatively evenly except within stem 1, which featured few mutations in the 5′ strand and many on the 3′ strand. The vast majority involved transitions (purine-purine or pyrimidine-pyrimidine conversions) rather than transversions. Most (62%) of the mutations in the stems also preserved the wild-type base-pairing by converting G:C pairs to G:U, A:U to G:U, or G:U to G:C. Finally, 5 of the 13 positions (Fig. 1 left, italic) that are protected from nuclease digestion in the SARS-CoV pseudoknot^4^ experienced mutation in SARS-CoV-2, close to the same rate of mutation as in the pseudoknot as a whole (38% vs. 48%), indicating that the protected residues are not significantly more resistant to undergoing mutation.

To examine the functional effects of these mutations on frameshifting, we selected a panel of 6 mutations (Fig. 1, magenta) located in regions that are important for the pseudoknot structure: near the stem 1/stem 2 and stem 1/stem 3 junctions (U20C, G29U, U58C), adjacent to an adenine bulge critical to frameshifting^4^ (C62U), and in the loop 2 dimerization domain^22^ (C43U, U47C). For each mutant, we measured the −1 PRF efficiency induced by the pseudoknot using cell-free translation of a dual-luciferase reporter system consisting of the *Renilla* luciferase gene in the 0 frame upstream of the firefly luciferase gene in the −1 frame and separated by the SARS-CoV-2 frameshift signal.^1^ The −1 PRF efficiency was obtained from the ratio of luminescence emitted by the two enzymes, compared to controls with 100% and 0% firefly luciferase read-through (SI Methods).

Comparing the results for each mutant to the −1 PRF efficiency seen for the consensus (wild-type) pseudoknot sequence, we found that 5 of the 6 mutations left the −1 PRF efficiency effectively unchanged within error (Fig. 2A). Mutations that disrupted base-pairing in a stem (*e*.*g*. G29U) or the dimerization domain (C43U, U47C) as well as for those that left all base-pairing intact (U58C, C62U) were both observed to have the same lack of effect, suggesting that most natural mutations do not alter the −1 PRF efficiency characteristic of the virus. Such a result is consistent with previous work suggesting that −1 PRF levels are regulated in a narrow range, outside of which the virus is significantly attenuated,^5–8^ and that they are not determined directly by characteristics such as stability or specific static structures but rather are most closely related to the conformational heterogeneity of the RNA.^23,24^ Remarkably, however, the U20C mutation, which disrupted the A:U base-pair at the end of stem 2 in the stem 1/stem 2 junction, caused a very significant decrease in −1 PRF efficiency, suppressing it over 3-fold (Fig. 2A, red). This result is particularly striking because in SARS-CoV, a ∼3.5-fold reduction of −1 PRF efficiency was shown to cause an over 1,000-fold attenuation of the virus.^5,6^ The fact that the U20C mutant was sampled from a COVID-19 patient indicates that SARS-CoV-2 can survive at a wider range of −1 PRF levels than expected based on work on other viruses.

**Fig. 2:**
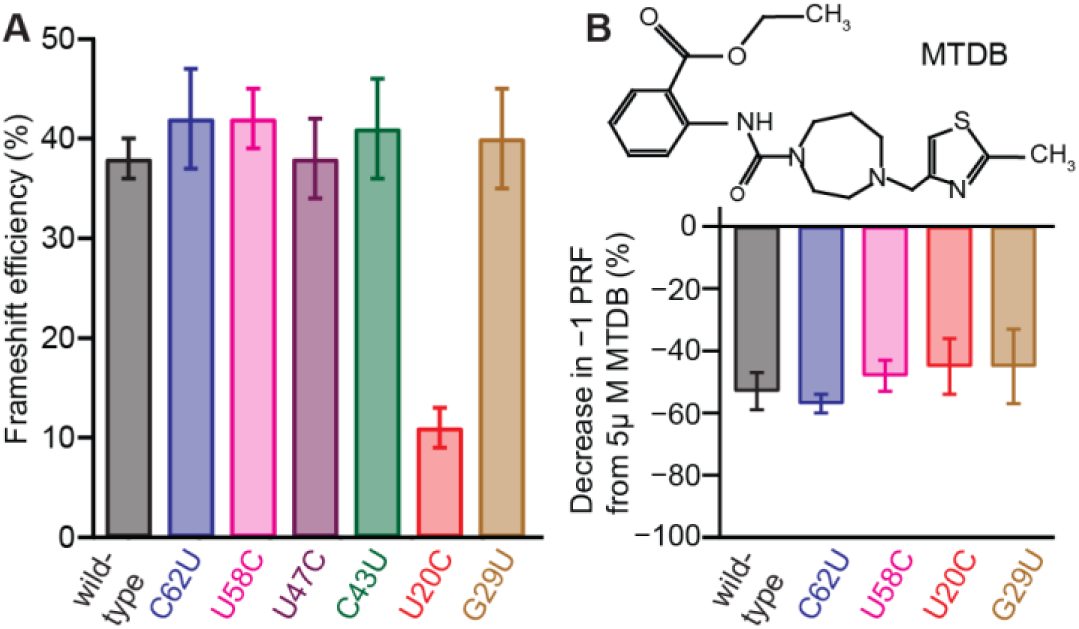
Effects of mutations on −1 PRF efficiency and anti-frameshifting activity of MTDB. (A) Most mutations left the −1 PRF efficiency unchanged, with the notable exception of U20C. (B) The anti-frameshifting activity of the small-molecule ligand MTDB was not affected by any of the mutations.

The surprising effect of the U20C mutation on −1 PRF may arise from a combination of key properties of U20 in the SARS-CoV-2 pseudoknot. First, this mutation destabilizes an inter-stem junction; stacking of paired bases across junctions is known to play an important role in the stimulatory power of pseudoknots.^25^ Structural modeling also indicates that U20 plays an important role in triplex-like interactions with loop 1 that organize the stem 1/stem 2 interface,^26^ and previous work showed that removing triples in pseudoknots can reduce −1 PRF efficiency.^27^ The relatively sparse network of triples predicted in this pseudoknot^26^ compared to others, in combination with the effects on the stem junction, could explain the sensitivity of −1 PRF to mutation of U20. Interestingly, G29U also disrupts an inter-stem junction, yet it leaves −1 PRF unchanged, suggesting that the stem 1/stem 3 interface is less important than the stem 1/stem 2 junction, consistent with work showing that stem 3 is not essential for frameshifting.^4^

Finally, we explored if these natural mutations might lead to resistance to the effects of an anti-frameshifting ligand, MTDB (Fig. 2B, inset), which was previously found to bind to the pseudoknot from SARS-CoV and inhibit −1 PRF.^9,11^ MTDB was recently shown to have a similar effect on −1 PRF in wild-type SARS-CoV-2.^1^ We repeated the dual-luciferase assays of −1 PRF for frameshift signals containing the G29U mutation in stem 1, U20C and C62U mutations in stem 2, and U58C mutation in stem 3 (omitting the mutations in loop 2, which are far from the proposed ligand binding site near the junction of the 3 stems^9^) while adding 5 μM MTDB, to quantify the reduction in −1 PRF caused by the ligand. Comparing to the effect of MTDB on −1 PRF when using the wild-type pseudoknot (Fig. 2B, black), we found that for all of the mutants, 5 μM MTDB reduced −1 PRF efficiency by roughly half. This result was obtained even for the U20C mutant, which already stimulated −1 PRF at a substantially lower efficiency. Hence none of mutations significantly altered the inhibition of −1 PRF by MTDB, regardless of whether the mutations altered the basal −1 PRF efficiency.

These results have important implications for therapeutic strategies targeting −1 PRF modulation. First, the example provided by U20C show that SARS-CoV-2 can clearly tolerate a substantial reduction in −1 PRF levels. Hence any putative drug suppressing −1 PRF may need to do so at quite dramatic levels to be effective therapeutically. In fact, MTDB itself induces a smaller decrease in −1 PRF than the U20C mutation and is thus unlikely to be effective as a drug for treating COVID-19. Just as important, however, is the evidence that the effects of an anti-frameshifting ligand can be insensitive to a wide range of natural mutations. The fact that the details of how the mutations affected the pseudoknot—whether they stabilized or destabilized the structure, disrupted base-pairing, affected one region of the structure or another, or even perturbed the pseudoknot enough to alter the basal −1 PRF rate significantly—were effectively immaterial to the activity of MTDB suggests that the inhibitory effect of the ligand is not easily evaded by simple mutations. Even though such an insensitivity to mutation was seen for only a single anti-frameshifting ligand, this behavior shows the promise of the pseudoknot as a therapeutic target, and holds out hope that a ligand with higher anti-frameshifting activity may be found in future work.

## Author contributions

KN and MTW designed the research; KN, SM, MZ, DBR, and SMI performed research; KN, SM, MZ, DBR, and MTW analyzed data; all authors wrote the manuscript.

## Acknowledgements

We gratefully acknowledge the originating laboratories that obtained SARS-CoV-2 specimens and submitting laboratories that generated and shared genetic sequence data via the GISAID Initiative, and all associated authors. This work was supported by the Canadian Institutes of Health Research and Alberta Innovates.

## Supporting Methods and Materials

### Preparation of mRNA constructs

A dual luciferase reporting system was created by cloning the sequence corresponding to *Renilla* luciferase and the multiple cloning site (MCS) from the plasmid pMLuc-1 (Novagen) upstream of the firefly luciferase sequence in the plasmid pISO (addgene), as described previously.^1^ The frameshift signal from SARS-CoV-2 was then cloned into MCS between the restriction sites *Pst*I and *Spe*I. Three different types of constructs were made. First, a construct for measuring −1 PRF stimulation was made, containing the frameshift signal with consensus (wild-type) slippery sequence (U UUA AAC) and consensus or mutant pseudoknot sequence, placing the downstream firefly luciferase gene in the −1 frame so that its expression was dependent on −1 PRF. Next, two controls were derived from this construct: (1) a negative control for measuring the background firefly luciferase luminescence (0% firefly luciferase read-through), in which the slippery sequence was mutated to include a stop codon (U UGA AAC); and (2) a positive control for measuring 100% firefly luciferase read-through, in which the slippery sequence was disrupted (U AGA AAC) and the firefly luciferase gene was shifted into the 0 frame. Sequences for all constructs are listed in Table S1.

Transcription templates were amplified from these plasmids by PCR, using a forward primer that contained the T7 polymerase sequence as a 5′ extension to the primer sequence.^1^ The mRNA for dual-luciferase measurements was produced from the transcription templates by *in-vitro* transcription (MEGAclear).

### Dual-luciferase frameshift assay

Frameshifting efficiency was measured by a cell-free dual-luciferase assay.^2^ Briefly, for each construct, 1 µg of mRNA transcript was heated to 65 °C for 3 min and then incubated on ice for 2 min. The mRNA was added to a solution mixture containing amino acids (10 µM Leu and Met, 20 µM all other amino acids), 35 µL of nuclease-treated rabbit reticulocyte lysate (Promega), 5 U RNase inhibitor (Invitrogen), and brought up to a reaction volume of 50 µL with water. The reaction mixture was incubated for 90 min at 30 °C. Luciferase luminescence was then measured using a microplate reader (Turner Biosystems). First, 20 µL of the reaction mixture was mixed with 100 µL of Dual-Glo Luciferase reagent (Promega) and incubated for 10 min before reading firefly luminescence, then 100 µL of Dual-Glo Stop and Glo reagent (Promega) was added to the mixture to quench firefly luminescence, and the reaction was incubated for 10 min before reading *Renilla* luminescence. The −1 PRF efficiency was calculated from the ratio of firefly to *Renilla* luminescence (F:R), subtracting the background F:R measured from the negative control and then normalizing by F:R measured from the positive control.

To quantify the effects of MTDB on −1 PRF, 5 μM MTDB was added to the reaction volume for each construct (wild-type and mutant pseudoknots, positive and negative controls). At least five replicates were measured and the results averaged, as described previously.^3^

**Table S1:** Sequences of measurement and control constructs. DNA sequence of constructs with slippery site (green) and linker region (blue). Pseudoknot mutation sites shown in bold.

|  |  |
| --- | --- |
| SARS-CoV-2 | T TTA AAC GGG TTT gcg gtg taa gtg cag ccc gtc tta cac cgt gcg gca cag gca cta gta<br>ctg atg tcg tat aca ggg ctt |
| C62U | T TTA AAC GGG TTT gcg gtg taa gtg cag ccc gtc tta cac cgt gcg gca cag gca cta gta<br>ctg atg tcg tat a**Ta** ggg ctt |
| U58C | T TTA AAC GGG TTT gcg gtg taa gtg cag ccc gtc tta cac cgt gcg gca cag gca cta gta<br>ctg atg tcg **C**at aca ggg ctt |
| U47C | T TTA AAC GGG TTT gcg gtg taa gtg cag ccc gtc tta cac cgt gcg gca cag gca cta g**Ca**<br>ctg atg tcg tat aca ggg ctt |
| C43U | T TTA AAC GGG TTT gcg gtg taa gtg cag ccc gtc tta cac cgt gcg gca cag gca **T**ta gta<br>ctg atg tcg tat aca ggg ctt |
| U20C | T TTA AAC GGG TTT gcg gtg taa gtg cag ccc g**C**c tta cac cgt gcg gca cag gca cta gta<br>ctg atg tcg tat aca ggg ctt |
| G29U | T TTA AAC GGG TTT gcg gtg taa gtg cag ccc gtc tta cac c**T**t gcg gca cag gca cta gta<br>ctg atg tcg tat aca ggg ctt |
| Positive control | T AGA AAC GGG TTT Tgc ggt gta agt gca gcc cgt ctt aca ccg tgc ggc aca ggc act agt<br>act gat gtc gta tac agg gct t |
| Negative control | T TGA AAC GGG TTT gcg gtg taa gtg cag ccc gtc tta cac cgt gcg gca cag gca cta gta<br>ctg atg tcg tat aca ggg ctt |

**Table S2: Accession numbers of mutant pseudoknot sequences in GISAID database.** Listed in file TableS2.xls

